# Predictions shape confidence in right inferior frontal gyrus

**DOI:** 10.1101/047126

**Authors:** Maxine T. Sherman, Anil K. Seth, Ryota Kanai

## Abstract

It is clear that prior expectations shape perceptual decision-making, yet their contribution to the construction of subjective decision confidence remains largely unexplored. We recorded fMRI data while participants made perceptual decisions and confidence judgements, controlling for potential confounds of attention. Results show that subjective confidence increases as perceptual prior expectations increasingly support the decision, and that this relationship is associated with BOLD activity in right inferior frontal gyrus (rIFG). Specifically, rIFG is sensitive to the discrepancy between expectation and decision (mismatch), and, crucially, higher mismatch responses are associated with lower decision confidence. Connectivity analyses revealed the source of the expectancy information to be bilateral orbitofrontal cortex (OFC) and the source of sensory signals to be intracalcarine sulcus. Altogether, our results indicate that predictive information is integrated into subjective confidence in rIFG, and reveal an occipital-frontal network that constructs confidence from top-down and bottom-up signals. This interpretation was further supported by exploratory findings that the white matter density of intracalcarine sulcus and OFC negatively predicted their respective contributions to the construction of confidence. Our findings advance our understanding of the neural basis of subjective perceptual processes by revealing an occipito-frontal functional network that integrates prior beliefs into the construction of confidence.

## Significance statement

Perceptual decision-making is typically conceived as an integration of bottom-up and top-down influences. However, perceptual decisions are accompanied by a sense of confidence. Confidence is an important facet of perceptual consciousness, yet remains poorly understood. Here we implicate right inferior frontal gyrus (rIFG) in constructing confidence from the discrepancy between perceptual judgement and its prior probability. Furthermore, we place rIFG within an occipito-frontal network, consisting of orbitofrontal cortex and intracalcarine sulcus, which represents and communicates relevant top-down and bottom-up signals. Together, our data reveal a role of frontal regions in the top-down processes enabling perceptual decisions to become available for conscious report.

Perception is increasingly being seen as an active process, in which current or future sensory states are inferred from predictive information (Engel et al., 2001; Lee, 2002; Bar, 2007; Beck and Kastner, 2009; Fiser et al., 2010; Gilbert and Li, 2013). These predictions can be modelled in Bayesian terms as prior beliefs, which bias perceptual inference towards solutions that are *a priori* more likely in a given context (Bülthoff et al., 1998; Seriès and Seitz, 2013; Trapp and Bar, 2015). Predictions, or priors, can have striking effects on perception, especially under high sensory uncertainty. For example, ambiguous rotational motion can be subjectively disambiguated by prior exposure to rotation direction, such that a mean rotation direction is perceived despite none existing in the physical stimulus (Maloney et al., 2005). In laboratory conditions, such behavioural effects of prediction are typically accompanied by increases in BOLD and ERP amplitude, as well as evoked gamma power, over sensory (Kouider et al. 2015; Egner et al. 2010; Saaltink et al. 2015; Kok et al. 2011; Jiang et al. 2013; Wacongne et al. 2011; Bauer et al. 2014) and decision-related (Bubic et al., 2009) brain regions - a ‘prediction error’ response profile that reflects the discrepancy between internal templates and perceptual content.

The perceptual content that forms the basis of our visual experience is accompanied by a degree of subjective confidence. Confidence reflects the estimated success of a perceptual choice, and can be seen as a gate for post-perceptual processes, such as learning and belief-updating (Nassar et al., 2010; Yeung and Summerfield, 2012). The communication of decision confidence can also facilitate group decision-making (Bahrami et al., 2010). Yet, while subjective confidence is an integral part of perceptual experience that can be easily probed in human subjects (Seth et al., 2008; Sandberg et al., 2010; Overgaard and Sandberg, 2012; Fleming and Lau, 2014; Wierzchoń et al., 2014), the construction of confidence remains poorly understood.

It is clear that confidence increases with evidence in support of the decision (Yeung and Summerfield, 2012; Hebart et al., 2014; Fetsch et al., 2015; Gherman and Philiastides, 2015). Decision and confidence are thought to evolve together until the first-order, objective decision has been made (Ratcliff and Starns, 2009; Kepecs and Mainen, 2012), and accordingly, there exists strong evidence for a common sensory signal underlying both types of report (Kiani and Shadlen, 2009; Fetsch et al., 2014; Kiani et al., 2014). Surprisingly, there has been much less research that considers the role of prior expectations on subjective confidence. There is converging behavioural evidence for subjective confidence increasing with prior evidence in favour of the associated choice (Aitchison et al., 2015; Meyniel et al., 2015a; Sherman et al., 2015), but the neural substrates of this have remained unexplored.

Here we aimed to identify brain regions in which prior perceptual expectations are integrated into confidence judgements. Based on previous work, we reasoned that confidence should be high when decisions are supported by prior knowledge, that is, when the discrepancy between expectation and perceptual decision is low. We therefore sought to identify brain regions that, first, are sensitive to both prediction error and confidence, and second, in which confidence is negatively associated with prediction error. In such a region, confidence would be associated with the mismatch between internal templates and perceptual report.

We further hypothesised that regions found to integrate prior expectations into confidence judgements (as described above) should be functionally connected with two information sources: one that represents the decision evidence, or sensory information; and one that represents the prior expectation. As confidence increasingly depends on prior expectations, functional connectivity with the source of the priors should increase. Similarly, when confidence is less dependent on priors, functional connectivity with the sensory region should increase.

## Materials and Methods

### Participants

The study was approved by the Brighton and Sussex Medical School Research Governance and Ethics Committee. Twenty-four healthy, English speaking and right-handed subjects were tested. Data from five participants were excluded: two whose thresholding failed (see section ‘Staircases’, Gabor hit rate = 2%, visual search d’ = −0.1); one for revealing abnormal vision only after scanning (and whose estimated contrast thresholds were accordingly > 2SD from the mean); one for excessive head movement in the scanner such that their T1 scan was unusable; and one for failing to respond on 33% of trials (relative to a mean of 3%). This left 19 participants with normal or corrected-to-normal vision for analysis. All participants gave informed, written consent and were reimbursed £50 for their time.

### Procedure

The experiment was conducted over three sessions at least 2 hours apart (no participant completed all three on a single day). In session one informed consent was obtained. Participants were trained on all tasks before scanning, which consisted of on-screen instructions, followed by a minimum of 10 practice trials of each task. Participants were encouraged to continue training until the task was well understood and response mappings learned.

To equate performance accuracy across conditions and subjects, participants subsequently completed three staircase procedures in the scanner but without acquiring echoplanar images (EPIs). Next, two 17 minute runs of experimental trials were completed while EPI scans were acquired.

Session two did not include a training component but was otherwise identical to session one. Session three consisted of: 10 minutes for T1 acquisition; 15 minutes of retinotopy (data from which is not used in this paper); and, time permitting, one more experimental run.

After three sessions participants were compensated for their time and debriefed.

### Experimental design

The paradigm used in the present study is adapted from a previously reported design(Sherman et al., 2015). The visual display was identical in all sections of the experiment (training, staircase and experimental). It consisted of a central visual search array and the presence or absence of a to-be-detected, Gabor patch in the periphery (see figure 1 and subsection ‘Trial Sequence’).

**Figure 1.**
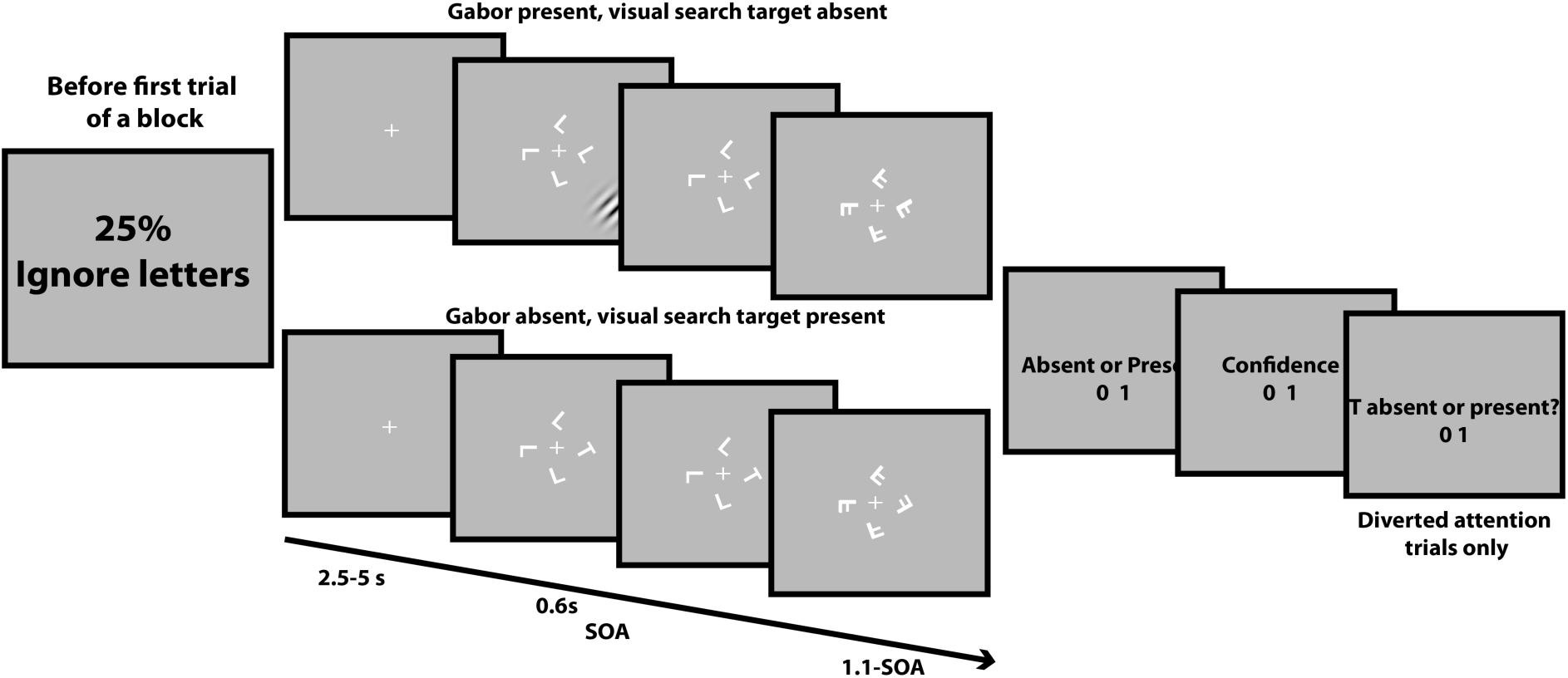
Trial sequence. Blocks began with instructions signalling the expectation and attention condition. On each trial a visual search target T was either absent (top) or present (bottom) with 50% probability. On each trial a target Gabor was either present (top) or absent (bottom) with probability determined according to condition. Response cues followed the offset of the stimuli. Staircase trials were identical, except there was no condition-specific instruction at the beginning and only task-relevant response cues were presented.

In experimental trials, the principal task was Gabor detection and two factors were orthogonally manipulated: prior expectations of Gabor presence and attention to Gabor detection. Expectations were manipulated block-wise, by changing the probability of target Gabor presentation (ℙ (Gabor present) = .25, .50 or .75). The ℙ(Gabor present) = .25 condition induced an expectation of Gabor absence, whereas the ℙ (Gabor present) = .75 condition induced an expectation of Gabor presence. The ℙ (Gabor present) = .50 condition acted as a control (flat prior). Attention was manipulated by instructing participants to either perform or ignore a visual search task presented concurrently to the Gabor target. This task consisted of detecting target ‘T’s amongst an array of distracter ‘L’s. Performing both tasks concurrently diverted attention from the Gabor detection task, allowing us to separate effects of expectation from those of attention.

These conditions were manipulated block-wise, in groups of 12 trials. Each condition occurred once per scanning run in fully counter-balanced order. Before each experimental block began participants were informed of both the expectation and attention condition via the presentation of an instruction screen presented for 10 seconds (see figure 1). Participants were instructed to always maintain fixation at a central cross.

### Trial sequence

The trial sequence was identical for training, staircasing and experimental trials and is shown in figure 1. Only instructions varied (see ‘Experimental design’). Trials began with a white fixation cross of random duration between 2.5 and 5 seconds. Next, a visual search array appeared, which consisted of seven letters: either all white, capital ‘L’s (50% chance), or a white, capital ‘T’ replacing an ‘L’ (50% chance). All letters were equidistant from fixation and took an independently random orientation. These were subsequently masked by a matching array of ‘F’s to increase task difficulty. In total the visual search array was present for 1.1 seconds. The stimulus onset asynchrony (SOA) between target and masking arrays was titrated for each participant such that accuracy was at 78% (see Staircases).

On some trials a near-threshold (see section Staircases) peripheral Gabor patch (orientation = 135°, phase 45° on 50% of trials, 225° on 50% of trials, sf = 2c/°, Gaussian SD = 30) was additionally presented. On these trials the stimulus appeared at the same time as the visual search array. To minimise attentional capture it was presented over 0.6 seconds in a Gaussian time envelope so that it had a gradual onset and offset. Stimulus contrast was titrated to equate performance across levels of attention and participants at 78% accuracy (see Staircases).

The interval between offset of the masking array and onset of response prompts was jittered during scanning only (i.e. experimental trials) to minimise motor cortex activity reflecting response anticipation. Jitter was randomly selected from the discrete values 1.3s:0.3s:3.1s.

Response prompts were presented at the end of the trial. The first prompt referred to the Gabor detection task. ‘Absent’ responses were recorded by pressing the outer left key and ‘present’ responses, the outer right key. This prompt was presented on all trials except those of the visual search staircase procedure (only visual search performed). The second prompt asked whether participants guessed (inner left) or were confident (inner right) in their Gabor detection response (not presented on staircasing trials). The third prompt was only presented on trials where participants performed the Gabor detection task and the visual search task together (dual-task trials). This asked whether the visual search target ‘T’ was absent (outer left) or present (outer right). Response prompts remained onscreen for 2 seconds and responses were coded as missed trials if no response was given within the allowed time.

### Staircases

Prior to each experimental session, three separate adaptive 1-up-3-down psychophysical staircase procedures (9 reversals) were completed in the scanner. Trials were identical to those in staircase trials (see Trial structure) except: there was no manipulation of attention or expectation; the Gabor was always present, but randomly oriented either 45° to the left or to the right; the Gabor task was 2AFC orientation discrimination instead of target detection; confidence ratings were not requested.

Staircase 1 titrated Gabor contrast to achieve 78% accuracy under full attention. Initial contrast was 1.5%. The visual search array was masked after 0.5 seconds. Participants were instructed to ignore the visual search array but still fixate centrally.

Staircase 2 titrated the SOA between the visual search array and masking array to set performance at 78% (in the visual search task). Initial SOA was 500ms. Participants ignored the 2AFC task and performed the visual search task. Here, the ignored Gabor was presented at the contrast acquired in staircase 1.

Staircase 3 titrated Gabor contrast to achieve 78% accuracy (in Gabor detection) under diverted attention. Initial contrast was set at that obtained in staircase 1 and visual search SOA was set at the value obtained by staircase 2. Here, participants performed both the Gabor and the visual search tasks. The visual search SOA was set at the value obtained in the previous staircase and initial contrast was set at that obtained in the first and titrated over the course of the staircase to obtain the diverted attention contrast level.

### Statistical analyses

Gabor detection sensitivity and decision threshold were quantified by computing type 1 signal detection theoretic (SDT) measures *d’* and *c* respectively. These are computed by classifying trials as hits (*h*), misses (*m*), false alarms (*fa*) or correct rejections (*cr*). Then,

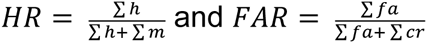

so that

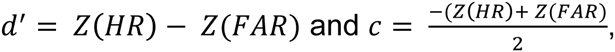

where Z is the inverse cdf of the normal distribution.

To obtain a measure of confidence threshold (i.e. bias towards reporting ‘confident’ or ‘guess’), which we denote as *C*, we used type 2 SDT. Here trials are classified as type 2 hits, misses, false alarms or correct rejections (Evans and Azzopardi, 2007). While in the type 1 case these are given by comparing stimulus class and response, in the type 2 case these are given by comparing decisional accuracy and confidence report. For example, a type 2 hit is a confident and correct response, whereas a type 2 false alarm is a confident but incorrect response. *C* is then defined analogously to *c*, but using type 2 hit rate and type 2 false alarm rate. This method allows us to quantify confidence relative to decision accuracy.

Behavioural and follow-up statistical tests were run on JASP (Love, J., Selker, R., Marsman, M., Jamil, T., Dropmann, D., Verhagen, A. J., Ly, A., Gronau, Q. F., Smira, M., Epskamp, S., Matzke, D., Wild, A., Rouder, J. N., Morey, R. D. & Wagenmakers, 2015). When the null hypothesis was predicted, Bayesian t-tests and repeated-measures ANOVAs implemented the JASP default Cauchy prior of 0.7 HWHM. All results presented were robust to reasonable adjustments of this value. Bayes factors greater than 1/3/10/100 are respectively interpreted as showing insensitive/moderate/strong/very strong evidence for the alternative hypothesis(Kass and Raftery, 1995). Bayes factors less than the reciprocal of these values are given the same labels, but refer to the null hypothesis.

Unless otherwise stated, all repeated-measures ANOVA results met the assumption of sphericity. Where sphericity was violated, corrected degrees of freedom and *p*-values are presented. The Greenhouse-Geisser correction is used for small violations (ε < .75) and the Huynh-Feldt correction for large violations (ε > .75).

### MRI acquisition and pre-processing

Functional T2* sensitive echoplanar images (EPIs) were acquired on a Siemens Avanto 1.5T scanner. Axial slices were tilted to minimise signal dropout from frontal and occipital cortices. 34 2mm slices with 1mm gaps were acquired (TR = 2863ms, TE = 50ms, FOV = 192mm x 192mm, Matrix = 64 x 64, Flip angle = 90°). Full brain T1-weighted structural scans were acquired on the same scanner and were composed of 176 1mm thick sagittal slices (TR = 2730ms, TE = 3.57ms, FOV = 224mm x 256mm, Matrix = 224 x 256, Flip angle = 7°) using the MPRAGE protocol.

Functional runs, each lasting 17 minutes, were collected per scanning session. Images were processed using SPM8 software (http://www.fil.ion.ucl.ac.uk/spm/software/spm8/). The first four functional volumes of each run were treated as dummy scans and discarded. Images were pre-processed using standard procedures: anatomical and functional images were reoriented to the anterior commissure; images were slice-time corrected with the middle slice used as the reference; EPIs were aligned to each other and co-registered to the structural scan by minimising normalised mutual information. Next, EPIs were spatially normalised to MNI space using parameters obtained from the segmentation of T1 images into grey and white matter. Finally, spatially normalised images were smoothed with a Gaussian smoothing kernel of 8mm FWHM.

### fMRI statistical analysis

At the participant level BOLD responses were time-locked to the onset of the visual search array (which appeared at the same time as the Gabor, if present), enabling us to examine BOLD responses to both target present and target absent trials. BOLD responses were modelled in a GLM with regressors and their corresponding temporal derivatives for each combination of the following factors: Attention (full, diverted), Expectation (25%, 50%, and 75%), Stimulus (target present, target absent), Report (yes, no) and Confidence (confident, guess). If a certain combination of factors had no associated trials for a particular participant, that regressor was removed from the participant’s first level model and contrast weights rescaled.

The reliability of the regression weights was maximised by entering data from all runs and sessions together, increasing the trial count per regressor. To avoid smearing artefacts, no band-pass filter was applied. Instead, low-frequency drifts were regressed out by entering white matter drift (averaged over the brain) as a nuisance regressor (Law et al., 2005). Nuisance regressors representing the experimental run and six head motion parameters were also included.

Comparisons of interest were tested by running one-sample *t*-tests against zero at the participant level, then running group-level paired *t-*tests on the one-sample maps. Unless otherwise stated, all contrasts at the group level were run with peak thresholds of *p* < .001 (uncorrected) and corrected for multiple comparisons at the cluster level using the FDR method.

We wanted to control for possible confounds between reaction speed and confidence (which correlate (Petrusic and Baranski, 2003; Grinband et al., 2006), and between individual or condition-wise differences in Gabor contrast and confidence (which correlate, Rahnev et al., 2011). To do this, a control GLM was computed. Here, each regressor was parametrically modulated by both Gabor contrast and reaction time. By design, in this model confidence was independent of reaction time and BOLD amplitude was independent of individual and condition-wise differences in stimulus contrast. The Results section reports analyses on our main model, i.e. that which does not model Gabor contrast and reaction speed. We did this because the control model has a four-fold increase in number of regressors, reducing statistical power. Nonetheless, all GLM analyses were replicated under our control model when using a peak threshold of *p* < .005. Crucially, all results under rIFG were also replicated when using a peak threshold of *p* < .001.

Functional ROIs were defined using the MarsBaR 0.42 toolbox (http://marsbar.sourceforge.net/download.html). Anatomical areas showing significant differences in BOLD were identified using the SPM Anatomy toolbox (Eickhoff et al., 2005) and Brodmann areas were identified using MRIcro (Rorden and Brett, 2000). Results of whole-brain analyses were plotted onto glass brains using MATcro (http://www.mccauslandcenter.sc.edu/CRNL/tools/surface-rendering-with-matlab).

### Psychophysiological interaction (PPI) analysis

The psychophysiological interaction analysis (PPI) was performed using the CONN functional connectivity toolbox (http://web.mit.edu/swg/software.htm). The GLM comprised regressors for attention condition (full/diverted), confidence (confident/guess) and expectation-response congruence (congruent/neutral/incongruent). Nuisance regressors were identical to those used in the GLM on BOLD. Again, the signal was not band-pass filtered but instead the mean WM drift was entered as a nuisance regressor. The data were denoised by regressing out signal from white matter, CSF and each individual condition, plus signal associated with all nuisance regressors. The PPI was run on univariate regression weights to identify effective connectivity between a functionally defined seed and remaining voxels. These weights were examined in a second level model which used an uncorrected peak threshold of *p* < .005 and FWE cluster corrected threshold of *p* < .05.

### Voxel-based morphometry (VBM)

T1-weighted structural scans were reoriented to the anterior commissure and segmented into grey matter (GM), white matter (WM) and CSF. These were normalised to MNI space using DARTEL with SPM defaults and a Gaussian smoothing kernel of 8mm FWHM (Ashburner and Friston, 2000). White matter and grey matter images were separately compared across participants in a multiple regression with age and total intracranial volume (GM + WM + CSF) as nuisance regressors. Including gender resulted in multicollinearity (older participants were more likely to be male) so gender was not modelled. Unless reported otherwise, clusters reported as significantly correlating with behaviour survived voxel-wise FWE correction.

## Results

### Expectations liberalise decisions and attention increases contrast sensitivity

Our first analyses confirmed the efficacy of our paradigm. To equate difficulty across attention conditions and participants, adaptive psychophysical staircases identified the stimulus contrast required for 78% accuracy on the Gabor detection task (see Methods subsection Staircases). Comparing the acquired contrasts in the full (*M* = 4.34%, *SD* = 3.50%) and diverted (*M* = 5.69%, *SD* = 3.79%) attention conditions revealed that contrast thresholds were significantly lower under full than diverted attention, *t*(19) = 2.95, *p* = .014, 95%CI [0.50%, 2.31%], d_z_ = 0.70 (fig. 2A). Thus, our paradigm successfully manipulated attention.

**Figure 2.**
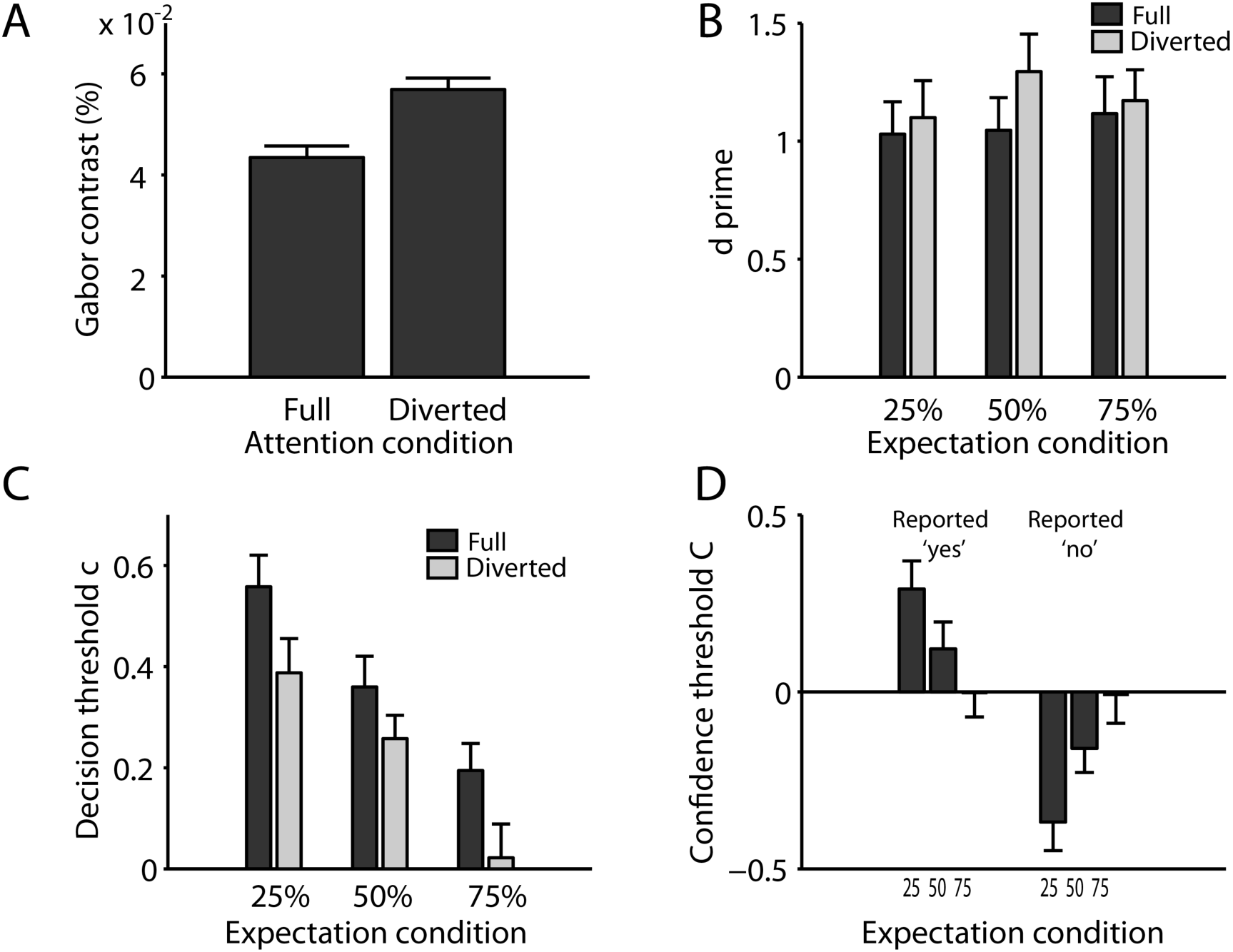
Behavioural effects of expectation and attention on objective and subjective decision-making. **A**. Stimulus contrast as a function of attention condition. To achieve 78% correct on the Gabor detection task contrast had to be higher under diverted than full attention. **B**. Detection sensitivity d’ as a function of expectation and attention condition. No significant differences were found. **C**. Decision threshold *c* as a function of expectation and attention condition. Independently of attention, bias towards reporting ‘yes’ (lower values of *c*) increases with the prior probability of Gabor presence **D**. Confidence threshold *C* as a function of expectation and perceptual report. Confidence for ‘yes’ responses increases (lower values of *C*) with increasing prior probability of target presence. Confidence for ‘no’ increases with increasing prior probability of target absence. Effect of expectation on type 2 C (confidence threshold). Therefore, confidence increases with expectation-response congruency. Error bars represent within-subjects SEM.

To ensure that our staircase procedure successfully equated detection sensitivity *d’* across conditions we ran a within-subjects Attention (full, diverted) × Expectation (25%, 50%, 75%) ANOVA. This revealed no significant difference between *d’* under full (M = 1.06, SE = 0.14) and diverted (M = 1.21, SE = 0.20) attention conditions, *F*(1,18) = 0.34, *p* = .569, η_p_^2^ = .02 (fig. 2B), and was corroborated by a Bayesian repeated-measures ANOVA of the same design that revealed moderate evidence for the null hypothesis (BF = 0.240). There was also no significant effect of Expectation on *d’*, *F*(2,36) = 0.70, *p* = .505, η_p_^2^ = .04, BF = 0.07 (strong evidence for the null) and no significant interaction term *F*(2,36) = 0.76, *p* = .476, η_p_^2^ = .04, BF = 0.016 (strong evidence for the null). Our staircases therefore successfully equated *d’*.

To determine whether we had successfully manipulated priors, we compared signal detection theoretic decision thresholds (*c*, see Methods) across expectation conditions (de Lange et al., 2013; Morales et al., 2015; Sherman et al., 2015). As the expectation of Gabor presence over absence increases, decision threshold should become increasingly biased towards ‘yes’ responses (i.e. liberalised, shown by smaller values of *c*). This was confirmed in a within-subjects Attention (full, diverted) × Expectation (25%, 50%, 75%) ANOVA, *F*(1.65, 29.72) = 18.10, *p* < .001, η_p_^2^ = .50. LSD post-hoc tests revealed a greater bias towards reporting ‘yes’ in the 50% (neutral) than the 25% (expect absent) condition, *p* = .010, d_z_ = 1.15, and greater still in the 75% (expect present) than the 50% (neutral) condition, *p* < .001, d_z_ = 1.39 (fig. 2C). We found no evidence for attentional effects on decision threshold, *F*(1, 18) = 3.38, *p* = .083, η_p_^2^ = .16, and no Expectation × Attention interaction, *F*(2, 36) = 0.37, *p* = .693, η_p_^2^ = .020. Summarising these results, our design successfully independently manipulated attention and expectation, while keeping detection sensitivity constant across conditions.

### Expectations liberalise confidence judgements

We have previously shown that subjective confidence increases when perceptual decisions are congruent with prior expectations (Sherman et al., 2015), and on this basis hypothesised that confidence would relate to prediction error signals. To determine whether we had replicated this behavioural result, we compared confidence for perceptual decisions that were congruent with expectations against those that were incongruent. Congruent responses are ‘yes’ reports in the 75% (expect present) condition and ‘no reports in the 25% (expect absent) condition. The reverse applies for incongruent responses. Confidence was analysed using type 2 SDT (see Methods for details). Broadly, while decision threshold *c* quantifies the extent to which perceptual decisions are biased towards ‘no’ reports, confidence threshold *C* quantifies bias towards reporting ‘guess’ over ‘confident’. An Attention × Expectation × Report ANOVA on confidence threshold *C* revealed that we replicated our previous finding. There was a significant Expectation × Report interaction, *F*(2,38) = 22.78, *p* < .001, η_p_^2^ = .555, such that when participants reported ‘yes’, participants became more likely to report decisions with confidence as the prior probability of target presence increased, *F*(1,18) = 17.00, *p* = .001, η^2^ = .486. Similarly, when participants reported ‘no’, high confidence became more likely as the prior probability of target absence increased, *F*(1,18) = 15.51, *p* = .001, η^2^ = .463 (figure 2D).

### Two forms of congruency

To unravel the neural correlates of predictive influences on confidence, we first needed to identify brain regions sensitive to perceptual expectations. We predicted, based on previous work, that areas sensitive to perceptual expectations would exhibit an increased BOLD amplitude for trials on which expectations were violated (Egner et al., 2010; Kok et al., 2011; Jiang et al., 2013; John-saaltink et al., 2015; Kouider et al., 2015). There are two possible ways to define expectancy violations here. Because the experimental design used near-threshold stimuli, leading to potential dissociations between percept and physical stimulus presentation, violations could occur with respect to either physical stimulus presentation, or perceptual report. We term the neural correlates of these types of incongruence PE_STIMULUS_ and PE_REPORT_ respectively. The former reflects the BOLD response to discrepancy between internal templates and stimulus presentation, whereas the latter reflects the BOLD response to discrepancy between internal templates and participants’ reported percept. PE_STIMULUS_ is most often observed at lower levels of the perceptual hierarchy^15–20^, whereas the decision-related PE_REPORT_ signals are often reported in higher-level, decision-related areas (Bubic et al., 2009), though they can be observed in visual cortex as well (Pajani et al., 2015).

### Representation of PE_STIMULUS_ in visual cortex

In our first analysis, we searched for regions that are sensitive to discrepancies between expectation and stimulus presentation (PE_STIMULUS_) over whole brain. To do this, we computed the contrast unexpected stimulus presentation > expected stimulus presentation. Target presence is expected in the 75% condition but unexpected in the 25% condition. Target absence is expected in the 25% condition but unexpected in the 75% condition. Our analysis identified one PE_STIMULUS_-sensitive area in contralateral occipital cortex (V1 to V3, BA18, peak MNI *x* = −12, *y* = −80, *z* = 22, Z_peak_ = 4.09, 0.66cm^3^, cluster *p*_FDR_ = .350, *p*_uncorr_ = .023) and one on the ipsilateral side (V1 to V3, BA18, peak MNI *x* = 8, *y* = −80, *z* = 18, Z_peak_ = 3.99, 1.01cm^3^, cluster *p*_FDR_ = .205, *p*_uncorr_ = .007). Neither of these clusters survived cluster-level correction, so they will not be considered beyond this point. They are presented to simply to show consistency with previous studies, in which statistical power was improved by constraining the analysis with functional localisers (Smith and Muckli, 2010; Kok et al., 2011, 2012; Larsson and Smith, 2012; Jiang et al., 2013) .

The whole-brain contrast PE_STIMULUS_, attended > PE_STIMULUS_, unattended yielded no significant or marginally significant clusters, indicating no evidence for a PE_STIMULUS_ × attention interaction.

Using a peak threshold of *p* < .005 both of these analyses were replicated under our control model, which included reaction speed and Gabor contrast as parametric modulators (unexpected > expected, contralateral: *p*_FDR_ = .446, *p*_uncorr_ = .014, ipsilateral: *p*_FDR_ = .446, *p*_uncorr_ = .011).

### Regions representing PE_REPORT_

Next, we searched for regions whose BOLD response reflects the discrepancy between expectation and perceptual report (PE_REPORT_). Expectation-congruent reports are ‘yes’ responses in the 75% (expect present) condition and ‘no’ responses in the 25% (expect absent) condition. The reverse applies for expectation-incongruent reports. These definitions differ from those in the previous analysis, because they consider perceptual report instead of stimulus presence or absence.

The contrast expectation-incongruent report > expectation-congruent report was computed over whole-brain. This revealed eight significant clusters reflecting PE_REPORT_, distributed throughout cortex (figure 3A and table 1). Our control analysis revealed that this effect was not driven by differences in Gabor contrast or reaction speed. We found no significant clusters for the reverse contrast, even with a more liberal peak threshold of *p* < .005 uncorrected.

**Figure 3.**
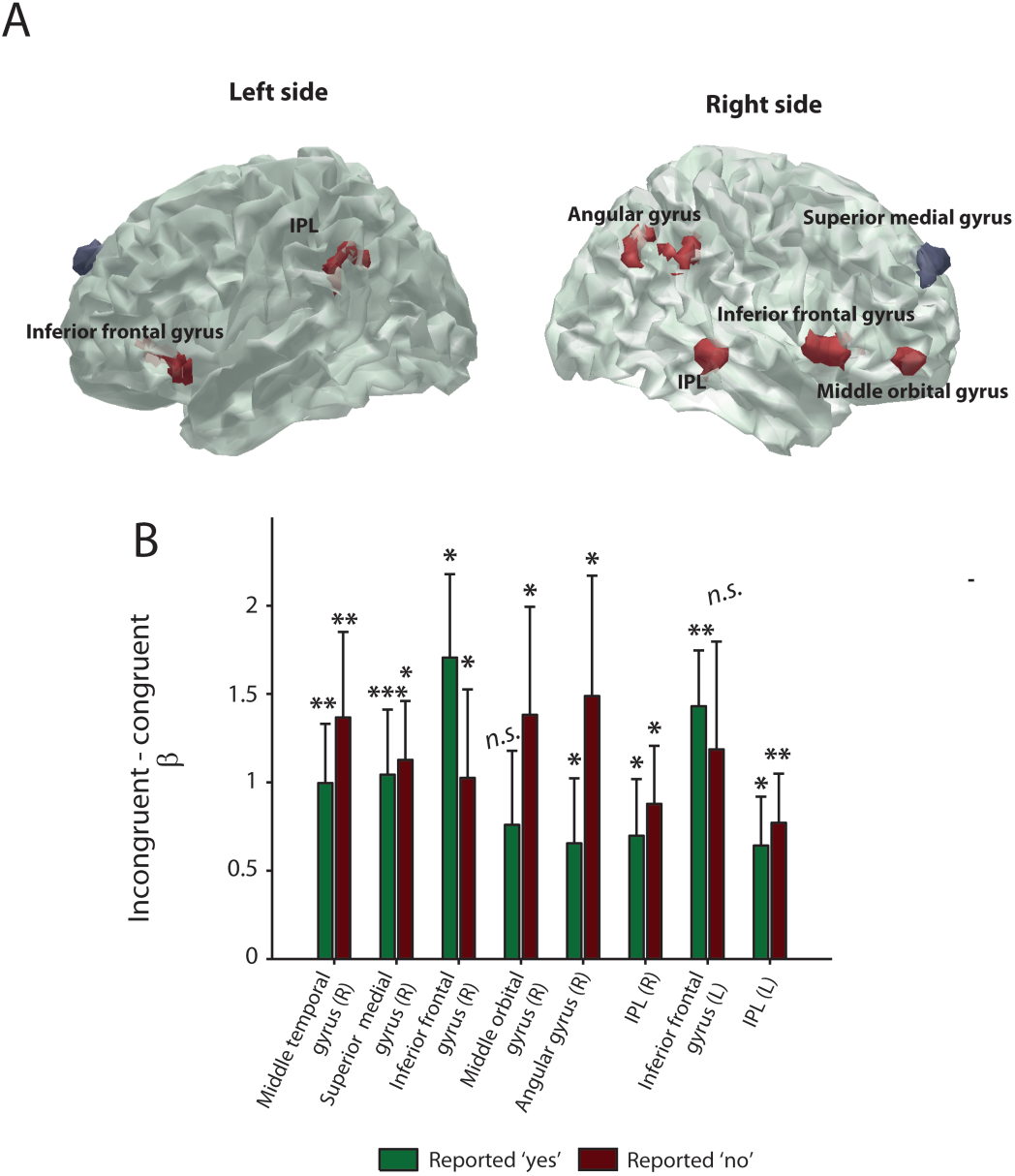
Report prediction error. **A**. Results of contrast incongruent response > congruent response over whole brain. Only clusters surviving FDR cluster-correction are shown. **B**. PE_REPORT_ (incongruent – congruent), by region and perceptual report. BOLD has been averaged over levels of attention. Stars represent whether PE_REPORT_ is significantly different from zero. Error bars represent SEM * *p* < .05, ** *p* < .01, *** *p* < .001

**Table 1.** Results of whole-brain analysis expectation-incongruent report > expectation-congruent report

| Region | BA | Side | Volume<br>(cm<sup>3</sup>) | Peak<br>z-value | $p_{FDR}$ | Peak MNI | | |
| --- | --- | --- | --- | --- | --- | --- | --- | --- |
|  |  |  |  |  |  | X | y | z |
| Middle temporal gyrus | 21 | R | 2.29 | 4.78 | .007 | 54 | -30 | -2 |
| Superior medial gyrus | 9/10 | R | 4.15 | 4.54 | < .001 | 12 | 58 | 32 |
| **Inferior frontal gyrus** | **47/48** | **R** | **2.70** | **4.45** | **.004** | **56** | **12** | **-2** |
| Middle orbital gyrus | 47/46 | R | 2.08 | 4.33 | .009 | 40 | 50 | -6 |
| Angular gyrus | 39 | R | 1.21 | 3.95 | .044 | 46 | -64 | 36 |
| Inferior parietal lobule | 40 | R | 1.21 | 3.91 | .044 | 58 | -40 | 40 |
| Inferior frontal gyrus | 47 | L | 1.90 | 3.79 | .012 | -38 | 26 | -4 |
| Inferior parietal lobule | 40/48 | L | 1.60 | 3.75 | .021 | -54 | -46 | 34 |

Regions exhibiting a PE_REPORT_ pattern should show heightened BOLD for incongruent responses irrespective of whether that response was a ‘yes’ or a ‘no’ (Kok et al., 2011). To test this in the above ROIs, median regression coefficients were extracted as a function of attention, expectation and report, and subjected to separate repeated-measures ANOVAs. Results are depicted in figure 3B and statistics are presented in table 2. All regions exhibited a significant PE_REPORT_ response for both ‘yes’ and ‘no’ judgements, except middle orbital gyrus and left inferior frontal gyrus. As a result these are not considered regions representing PE_REPORT_.

**Table 2.** Effect of expectation, separately for ‘yes’ and ‘no’ reports. Both effects should be significant for the region to be deemed a PE_REPORT_ region

| Region | Reported ‘no’ |  |  | Reported ‘yes’ |  |  | PE<sub>REPORT</sub><br>represented |
| --- | --- | --- | --- | --- | --- | --- | --- |
| | *F* | *p* | $\eta^2$ | *F* | *p* | $\eta^2$ | |
| Middle temporal gyrus | 8.82 | .008 | .329 | 5.83 | .006 | .245 | Yes |
| Superior medial gyrus | 8.10 | .001 | .310 | 4.46 | .014 | .213 | Yes |
| **Inferior frontal gyrus**<br>**(R)** | **4.70** | **.015** | **.207** | **3.45** | **.041** | **.162** | **Yes** |
| Middle orbital gyrus | 1.95 | .157 | .098 | 3.42 | .044 | .160 | No |
| Angular gyrus | 3.52 | .040 | .164 | 4.07 | .025 | .185 | Yes |
| Inferior parietal lobule<br>(R) | 4.71 | .044 | .207 | 7.17 | .015 | .285 | Yes |
| Inferior frontal gyrus (L) | 5.62 | .008 | .238 | 2.87 | .070 | .137 | No |
| Inferior parietal lobule<br>(L) | 5.39 | .032 | .230 | 6.04 | .005 | .251 | Yes |

All significant results here were replicated (at least at marginal significance) under our control model (for rIFG, our critical region, *p*_FDR_ = .044). Results were fully replicated when using a peak threshold of *p* < .005.

We have therefore identified six regions signalling PE_REPORT_: Right middle temporal gyrus (rMTG); right superior medial gyrus (rSMG), right inferior frontal gyrus (rIFG); right angular gyrus (rAG); and bilateral inferior parietal lobule (lPL). These results implicate this set of regions as having sensitivity to the discrepancy between perceptual expectations and perceptual choice.

### High confidence is associated with an attenuated PE_REPORT_ response in right IFG

Our main hypothesis was that high confidence would be associated with low PE_REPORT_. However, confidence can be also influenced by attention (Rahnev et al., 2011) and tracks accuracy (Dienes, 2008; Pleskaca nd Busemeyer, 2011). To test whether any PE_REPORT_ region represented confidence after controlling for these potential confounds, median regression weights from each PE_REPORT_ region were extracted as a function of confidence, attention and decision accuracy. These regression coefficients were then subjected to separate Bayesian repeated-measures ANOVAs. We were looking for regions whose BOLD response (in these regions, representing PE_REPORT_) differs with confidence. Note that we could not test for a PE_REPORT_ × Confidence interaction because the participant has signalled low confidence yes/no decisions as unreliable, that is, their perception of Gabor presence or absent does not necessarily correspond to their report.

Only one region demonstrated a BOLD response (i.e. PE_REPORT_ amplitude) that differed as a function of subjective confidence: rIFG. Here, supporting our hypothesis, BOLD amplitude was higher for guess than confident reports (figure 4A). Crucially, the analysis revealed substantially more evidence for modelling rIFG BOLD as a function of confidence alone (BF = 13.620) than as a function of just accuracy (BF = 0.877), just attention (BF = 0.711), or as a combination of confidence and any other factors (BF = 0.003 - 2.069, see table 3 for summary of results from all ROIs). A frequentist ANOVA gave the same result: a significantly higher BOLD amplitude for guess than confident responses, *F*(1,18) = 6.04, *p* = .024, η^2^ = .251, 95% CI [0.10, 1.28]. These results are depicted in figure 4B.

**Figure 4.**
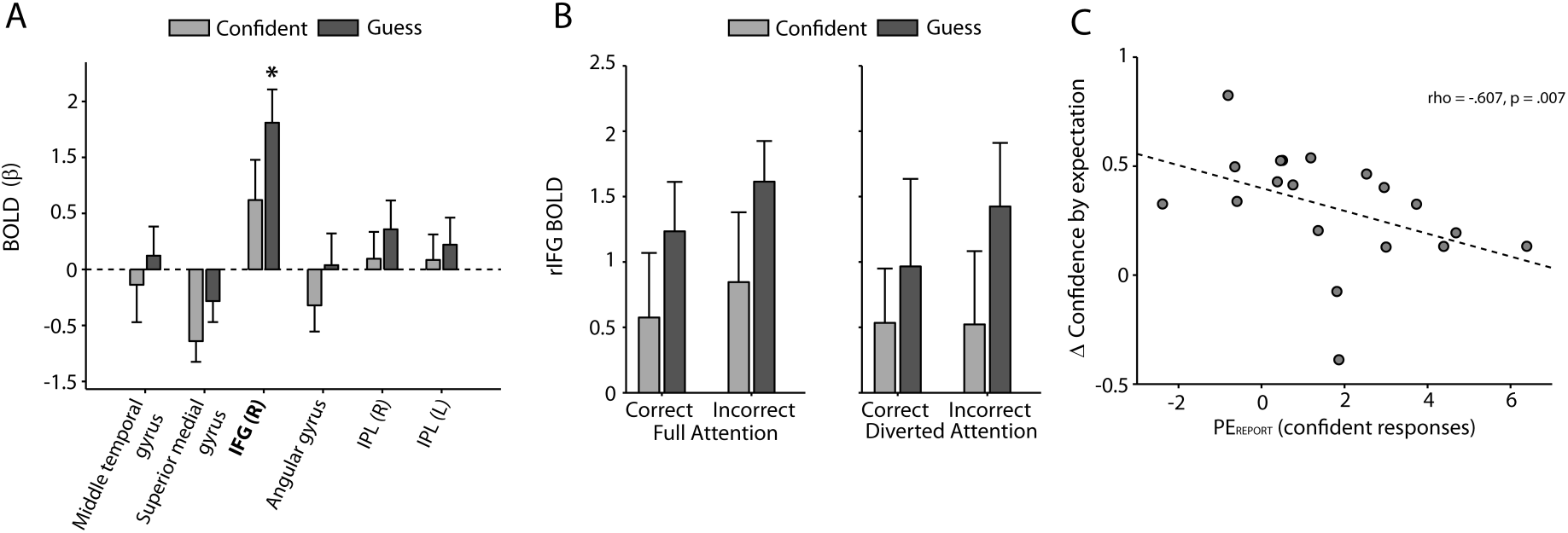
The relationship between confidence and report prediction error. **A**. BOLD as a function of confidence in each PE_REPORT_ region. BOLD is higher for guess than confident responses in rIFG only. **B**. rIFG BOLD is higher for guess than confident responses independently of attention and decision accuracy **C**. Brain-behaviour correlation. The higher the PE_REPORT_ response (confident reports only), the less that expectations increased confidence. Error bars represent +/- SEM. *\* p* < .05, ** *p* < .01 *** *p* < .001

**Table 3.**
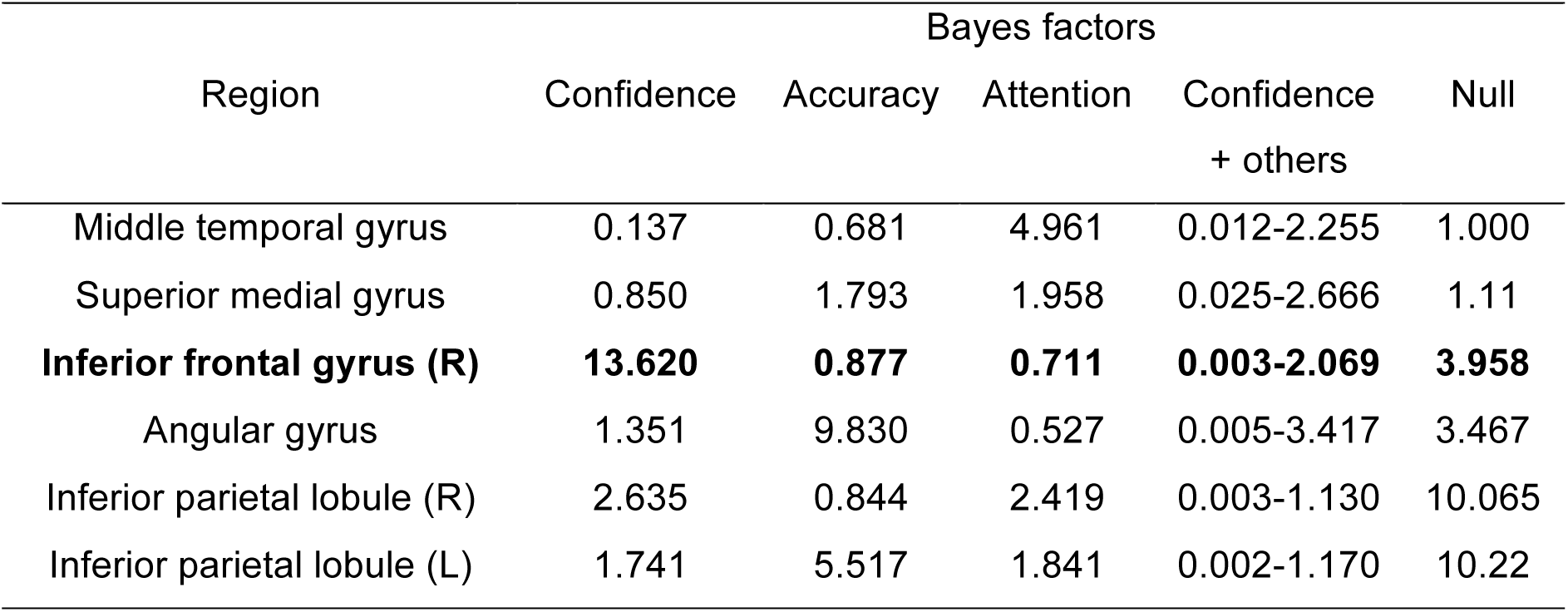
Results of Bayesian Confidence x Accuracy x Attention repeated-measures ANOVAS. Bayes factors correspond to the evidence for the listed model relative to the evidence for all other models

Next, we wanted to confirm that the effect of confidence on rIFG BOLD indeed reflects changes in PE_REPORT_. To do this, we restricted our analysis to confident responses and asked whether PE_REPORT_ decreases as expectations exert stronger influences on behavioural confidence. This would show that high confidence is associated with low PE_REPORT_ amplitude (i.e. a low expectation-report mismatch response). Furthermore, it would show that our behavioral effect of expectation on confidence is reflected in rIFG BOLD.

To test this, we calculated Δ*C* = incongruent type 2 *C* - congruent type 2 *C*. The group-level mean of Δ*C* is reflected in the slopes of figure 2D. This quantity reflects the extent to which confidence judgements are influenced (or weighted) by expectations. Next, we computed the BOLD difference between incongruent and congruent reports (PE_REPORT_), restricted to confident responses. Results showed that these quantities were negatively correlated, ρ = -.607, *p* = .007 (fig. 4C), confirming our finding that high confidence is associated with low PE_REPORT_ in rIFG: the more expectation increased confidence behaviourally, the more confidence was associated with low rIFG PE_REPORT_.

To ensure that these differences were not driven by differences in reaction speed or Gabor contrast, we extracted data from the cluster revealed by our control GLM. This revealed that even after controlling for these possible confounds, rIFG BOLD was significantly higher for guess that confident responses *t*(18) = 2.21, *p* = .041, d_z_ = 0.44. The significant brain-behaviour correlation was also replicated, rho = -.575, *p* = .014.

Together, these analyses reveal that subjective confidence is reliably associated with PE_REPORT_ in right IFG, even after controlling for attention, Gabor contrast, decision accuracy and reaction speed.

### Sources of priors and sensory signals for confidence

We have shown that rIFG activity associates response prediction error with confidence. Assuming a model in which decision confidence is a weighted function of top-down expectations and ‘bottom-up’ sensory signals (or decision evidence), we asked whether we could identify sources of these variables. To do this we ran a seed-to-voxel psychophysiological interaction analysis (PPI), with rIFG as a functionally defined seed.

We were interested in regions communicating predictive information, and therefore regions of interest would demonstrate functional connectivity with rIFG that differs for congruent and incongruent reports. We reasoned that while confidence should be a function of both sensory signals and expectations, there would be individual differences in how each component would be weighted, reflecting, for example, how reliable the expectation information is thought to be. Capitalising on these individual differences, we reasoned that rIFG would show stronger functional connectivity with the expectation region in participants whose confidence was weighted more by expectation. By contrast, rIFG would show stronger functional connectivity with the source of sensory signals in participants whose confidence was only weakly shaped by expectation.

To test this hypothesis we used a behavioural covariate of interest – the influence of expectations on confidence. This weighting function was assumed to be a function of the aforementioned behavioural variable Δ*C* = *C*_Incongruent_ – *C*_Congruent_., where *C* denotes confidence thresholds. This quantity is the same as the behavioural variable in figure 4B. Higher values signify that expectations exerted a stronger influence on confidence.

Sources of predictive information for confidence were identified by computing the contrast incongruent ≠ congruent, with Δ*C* as a between-subjects covariate of interest.

As shown in figure 5 (A-C), the PPI analysis revealed three significant clusters. The more expectations shaped confidence (higher Δ*C*), the more that congruence was associated with functional connectivity (FC) between rIFG and left orbitofrontal cortex (Δ*C* x congruent > incongruent; peak MNI *x* = −34, *y* = 36, *z* = 20, 2.82cm^3^, cluster P_FDR_ = .020) and right orbitofrontal cortex (Δ*C* x congruent > incongruent; peak MNI *x* = 10, *y* = 26, *z* = −18, 2.16cm^3^, cluster P_FDR_ = .036). On the other hand, the less expectations shaped confidence the more that congruence was associated with FC between rIFG and intracalcarine sulcus (Δ*C* x congruent > incongruent: peak MNI *x* = 6, *y* = −58, *z* = 12, 2.92cm^3^, cluster P_FDR_ = .025). Thus, intracalcarine sulcus and bilateral orbitofrontal cortices (OFC) exhibited a push-pull relationship, with the dominant region predicted by Δ*C*.

**Figure 5.**
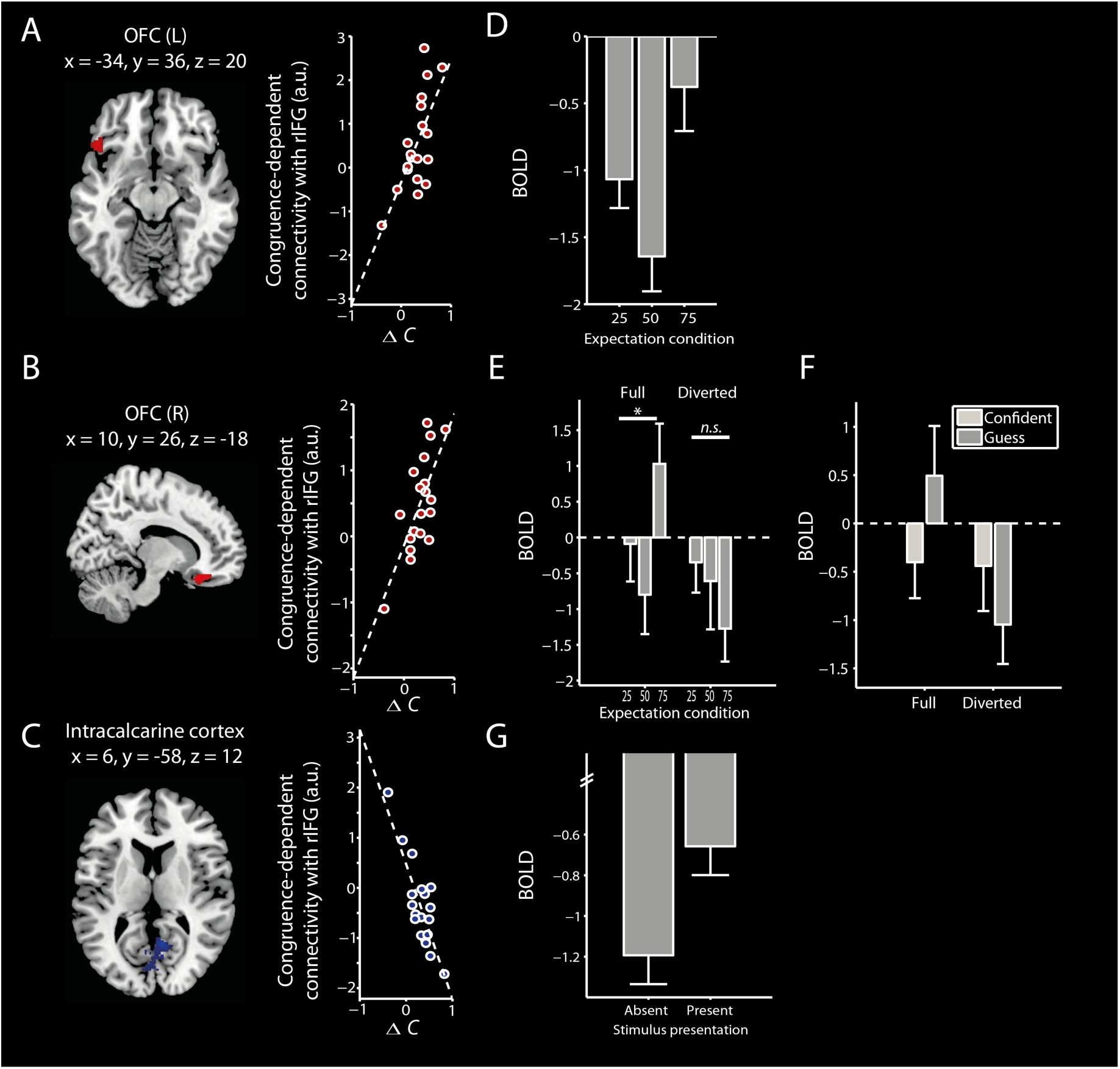
Results of PPI analysis, depicting functional connectivity with rIFG that differs for congruent and incongruent reports and scales with Δ*C*. Higher values of Δ*C* were associated with greater congruency-dependent connectivity with left **(A)** and right **(B)** orbitofrontal cortex, and less congruency-dependent connectivity with intracalcarine sulcus **(C)**. Scatterplots to the left of each cluster depict the correlation between Δ*C* and congruence-dependent functional connectivity. Panels **D-F** depict results from follow-up analyses in these regions. **(D)** Left OFC BOLD as a function of expectation. **(E)** Right OFC BOLD as a function of expectation and attention. **(F)** Right OFC BOLD as a function of confidence and attention. **(F)** Intracalcarine sulcus BOLD as a Gabor presence and absence. This difference is only marginally significant. Error bars represent within-subjects SEM.

Although the *balance* of FC between these regions was determined by Δ*C*, FC between these rIFG and these regions was present independently of Δ*C*. Specifically, FC between rIFG and OFC was significantly stronger on congruent than incongruent trials (lOFC *p* = .009, rOFC *p* = .043), whereas intracalcarine sulcus-rIFG FC was marginally stronger on incongruent than congruent trials (*p* = .078).

Because bilateral OFC was primarily associated with congruent responses, we reasoned that functional connectivity with these regions might reflect the communication of perceptual priors. Consistent with this, we found a main effect of expectation condition on lOFC BOLD *F*(2,36) = 3.61, *p* = .037, η_p_^2^ = .167, such that the expectation and BOLD were related by a ‘U’ shape function (fig. 5D), characteristic of the representation of prior information. This ‘U’ represents prior information because BOLD is higher when there is an informative prior (the 25% and 75% conditions) than when the prior is flat, or neutral (the 50% condition). In rOFC, this pattern was exhibited under full (*F*(2,36) = 3.51, *p* = .040, η_p_^2^ = .163, but not diverted, *F*(2,36) = 1.04, *p* = .363, η_p_^2^ = 0.55, attention (interaction *p* = .030, fig. 5E). Interestingly, in rOFC there was also a significant attention by confidence interaction, *F*(1,18) = 7.87, *p* = .012, η_p_^2^ = .304 (fig. 5F), such that attention reversed the BOLD response to confident versus guess responses.

These results are consistent with the interpretation of bilateral OFC communicating prior information. While lOFC represented prior information independently of attention, rOFC did this only under full attention. Moreover, the attention by confidence interaction under rOFC BOLD suggests that this region may additionally represent the degree of (reverse) uncertainty associated with attentional state.

We next asked whether intracalcarine sulcus represented prediction error signals. Response prediction error is demonstrated in an expectation by report interaction, whereas stimulus prediction error is demonstrated in an expectation by stimulus interaction. However, neither analysis was significant (both *p* > .441). Rather, the BOLD response here was marginally higher for stimulus present than absent trials, *F*(1,18) = 3.53, *p* = .077, η_p_^2^ = .164 (fig. 5G).

One might wonder whether bilateral OFC directly signals priors to intracalcarine sulcus, or vice versa for sensory signals. This was not the case. Re-running the PPI analysis in the same way, but with each OFC cluster as our seed revealed no significant or marginally significant connectivity with intracalcarine sulcus. Similarly, running the analysis setting each intracalcarine sulcus as the seed revealed no significant or marginally significant connectivity with either OFC cluster.

Taken together, these results suggest that the predictive information signalled to (or from) rIFG is a balance of priors and sensory signals. Moreover, they suggest that expectation-induced biases in subjective confidence are associated with increased rIFG-OFC functional connectivity, relative to sensory regions.

### The contribution of visual regions and OFC to confidence is predicted by white matter density

Our connectivity analyses revealed that OFC and intracalcarine sulcus represent priors and sensory signals respectively, and that the balance of rIFG connectivity with these regions differed across individuals. The presence of these individual differences motivated an exploratory follow-up analysis that asked whether they are reflected in brain structure. More specifically, we considered whether the weighting of top-down predictions and bottom-up prediction errors was a function of white or grey matter (WM and GM respectively) density of the source regions (i.e. OFC and intracalcarine sulcus).

The BOLD response of our cluster in OFC reflected an effect of perceptual expectations on objective decision. The behavioural correlate of this is therefore Δ *c* = *c*_*25*%_ - *c*_*75*%_, - the extent to which perceptual expectations bias (yes/no) decision. We performed a whole-brain multiple regression analysis on WM density, with total intracranial volume and participant age as nuisance covariates, and with Δ*c* as the regressor of interest. This analysis revealed that propensity to incorporate low-level priors into decision-making, as measured by Δ*c*, was negatively correlated with rOFC white matter density (fig 6A and B, peak MNI *x* = 23, *y* = 30, *z* = −14, 11.51cm^3^, P_peak-FWE_ = .030, Z = 5.08). The same analysis for GM yielded no significant results.

**Figure 6.**
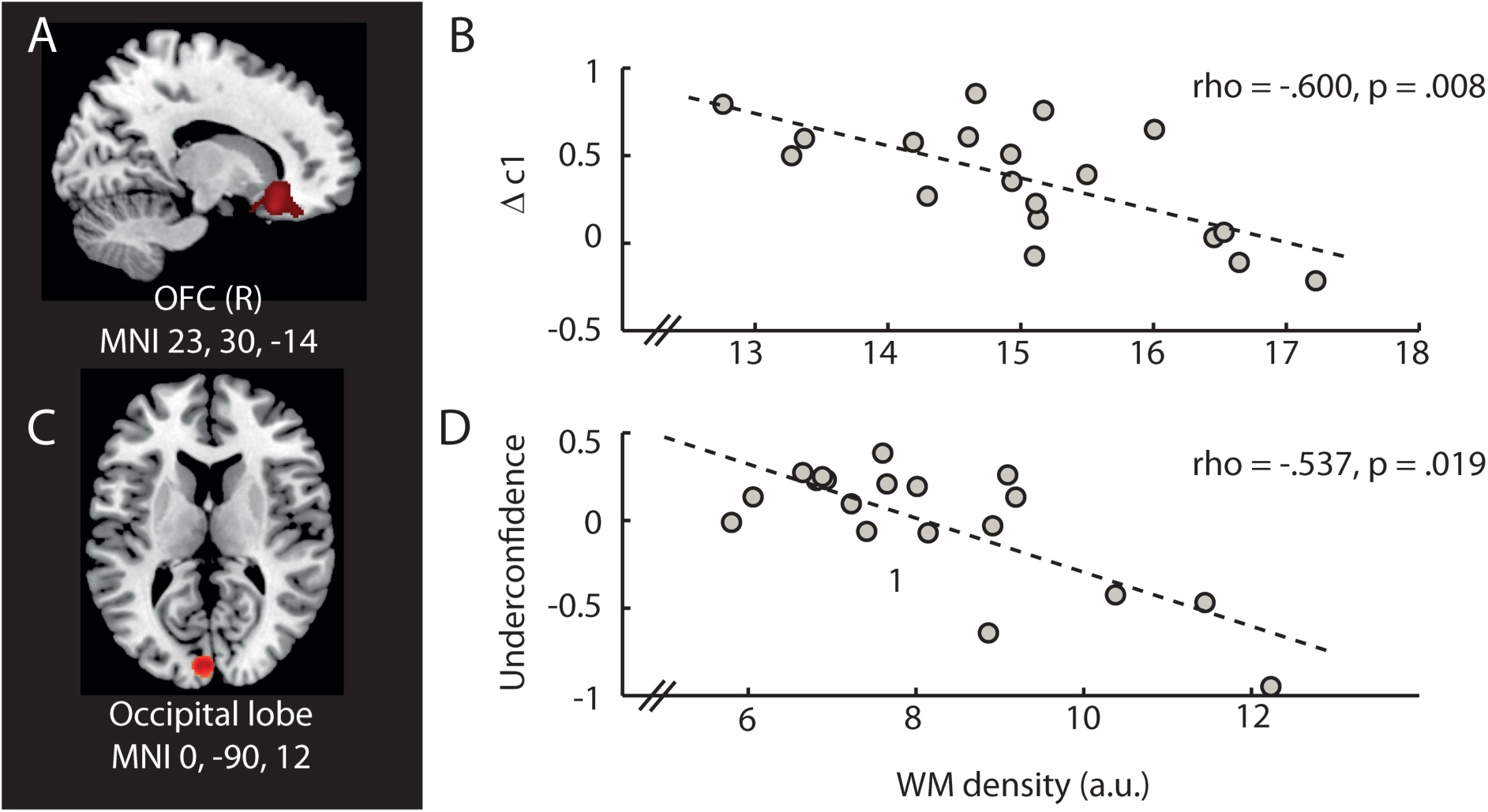
VBM results. **A, B**. White matter density in right orbitofrontal cortex negatively predicts the effect of expectation on perceptual decision. **C,D**. White matter density in contralateral occipital pole is negatively correlated with underconfidence bias (mean confidence threshold).

Given that both rOFC and rIFG BOLD predicted confidence we performed the same analysis, but this time with mean confidence threshold as the regressor of interest. Confidence threshold represents one’s overall belief in their perceptual performance, such that lower values as associated with higher confidence in decision-making ability. This revealed a significant cluster in contralateral occipital lobe. Here, increasing WM density significantly predicted more liberal confidence thresholds at the cluster, but not the peak level (figure 6C and D peak MNI *x* = 0, *y* = −90, *z* = 12, 6.31cm^3^, P_peak-FWE_ = .879, P_FDR_ = .031).

Together these results suggest that the dependence of confidence on functional connectivity with source regions is reflected in anatomical indications of that connectivity: WM density in OFC was negatively predicted by its functional correlate; and increasing occipital pole WM density was associated with mean confidence, that is, beliefs of better perceptual performance.

## Discussion

In the present paper we have shown that behavioural confidence in perceptual decision increases when decisions are supported by (or congruent with) prior expectations. Crucially, we show that this predictive information is, at least in part, integrated into confidence in right inferior frontal gyrus (rIFG).

We have shown that unexpected percepts, taken with respect to the decision or report, are associated with a heightened BOLD response (termed here PE_REPORT_) in a distributed set of frontal, parietal and temporal decision-related regions. Interestingly, this expectation-sensitive set resembles those implicated in other forms of ‘top-down’ processing such as modality-independent sensory change detection (Downar et al., 2000), response inhibition (Verbruggen and Logan, 2008; Criaud and Boulinguez, 2013), and detection of behavioural salience (Downar et al., 2002).

Our crucial result was that the contribution of top-down expectations to subjective confidence judgements was reflected in fMRI BOLD, specifically in right inferior frontal gyrus (rIFG). Here, high confidence was associated with a lower prediction error response profile. Furthermore, the more that confidence was shaped by expectation behaviourally, the more that confidence was associated with low prediction error signals in this area. Our results therefore indicate a central role for rIFG in perceptual decision making in which the ‘match’ between internal templates and perceptual content is integrated into subjective confidence judgements.

Under an alternative account, the sensitivity of rIFG to confidence would be an indirect effect of sensitivity to task difficulty. For example, rIFG may infer task difficulty from the degree to which the percept is surprising. However, this interpretation was ruled out by control analyses, which showed that the PE_REPORT_-confidence relationship was not driven by choice accuracy. These control analyses additionally excluded attention, stimulus contrast, and reaction speed as driving the observed relationship between PE_REPORT_ and confidence in rIFG.

This process of relating predictive information into confidence judgements recruited both intracalcarine sulcus, primarily when incongruent reports were made, and bilateral orbitofrontal cortices, primarily when congruent reports were made. The former region showed a marginally significant BOLD response to the stimulus. We assume that this effect was weak because the effect was localised to a large cluster, while reflecting the neural response to a small stimulus in retinotopically-organised space. We interpret the functional connectivity with intracalcarine sulcus as the communication of sensory signals. Broadly in this region, white matter density predicted participants’ overall level of confidence (confidence threshold), implicating this perceptual region in shaping beliefs in one’s perceptual performance. By contrast, we found that both right and left orbitofrontal cortex represented prior information, consistent with previous work (Schoenbaum and Roesch, 2005; Wallis, 2007; Trapp and Bar, 2015). Interestingly, the representation of attentional state was lateralized in these areas: left OFC represented prior information independently of attention, whereas right OFC was sensitive to attentional state. Here, representation of the prior required attention, and furthermore, the BOLD response to decision confidence reversed with attention. Under full attention rOFC BOLD was higher for guess responses than confident responses, as is usually found (Fleming et al., 2012; Hilgenstock et al., 2014). However, under diverted attention this pattern reversed, possibly indicating that rOFC represents the uncertainty associated with attentional state: high under full attention, but low under diverted attention. Increasing white matter density in rOFC was associated with the behavioural effect of expectations on decision, supporting our interpretation of its role in representing the prior.

Altogether, we interpret these results as showing that subjective confidence is represented in rIFG as a combination of both stimulus-driven signals, communicated from intracalcarine sulcus, and shaped by top-down perceptual expectations, communicated from bilateral OFC. OFC has been repeatedly been shown to reflect reward expectations and beliefs (Kepecs et al., 2008; Kim et al., 2011; De Martino et al., 2013; Lebreton et al., 2015), however here we place OFC belief representations within a larger hierarchical structure for perceptual processing, generating predictions (Stalnaker et al., 2015; Trapp and Bar, 2015) that constrain subjective confidence judgements in perceptual decision. Importantly, our PPI analysis cannot determine the directionality of functional connections in this network. One possibility is that rIFG is involved in constructing confidence from an integration of PE_REPORT_ signals and top-down expectations. Here, both intracalcarine sulcus and OFC would be sending signals to rIFG. However, another possibility is that PE_REPORT_ signals are passed from occipital lobe to rIFG, and an initial transformation of PE_REPORT_ into confidence is signalled to rOFC. Under such an account, the role of rOFC here may be one which transforms the confidence estimate represented in rIFG into a reportable judgement, based on the mismatch between the estimate, expectations, and potentially, attentional state (Lebreton et al., 2015). Further studies will be needed to disambiguate these possibilities.

Our results are readily interpretable from Bayesian brain perspectives (Lee, 2002; Yuille and Kersten, 2006; Friston, 2009; Clark, 2013). These propose that perceptual inference is a weighted integration of sensory evidence and prior beliefs about the cause of the sensation, such that the perceptual report corresponds to the belief with the greatest posterior probability. The posterior probability increases as the correspondence between prior and sensory signal increases. Therefore, inference is deemed ‘successful’, and so should be associated with high confidence, when we see a low ‘prediction error’ response, as we saw here (Meyniel et al., 2015b). Neuronal representations of prediction errors are well-established in the reward domain (Nakahara et al., 2004; Bayer and Glimcher, 2005), but in the perceptual domain evidence remains restricted to BOLD correlates such as PE_REPORT_. Under such a Bayesian brain account, our connectivity results suggest that intracalcarine sulcus passes sensory signals to rIFG, and rOFC passing top-down predictions. In this view, the finding that PE_REPORT_ amplitude in rIFG was lower for confident responses is consistent with the representation or construction of the posterior belief in this region. This in turn is in line with empirical evidence for rIFG encoding of the decision variable, either in Bayesian form (the posterior; d’Acremont et al. 2013) or as decision evidence (Hebart et al., 2014), which are mathematically equivalent, Bitzer et al. 2014).

Previous work has separately implicated rIFG in the representation of both the decision variable (Bubic et al., 2009; d’Acremont et al., 2013) and expectation violation in a range of modalities, from speech perception (Clos et al., 2014) to auditory deviance detection (Garrido et al., 2009) and visual perception (Bubic et al., 2009). Previous work has also implicated rIFG in the representation of subjective uncertainty (Fleck et al., 2006; Fleming and Dolan, 2012). However, to our knowledge these variables had not been related to each other before. rIFG has also been implicated in a wide range of related executive processes, such novelty detection (Hampshire et al., 2010), change detection (Beck et al., 2001), and behavioural relevance (Hampshire et al., 2010), including, crucially, detecting or resolving response conflict (Casey et al., 2000; Hampshire et al., 2010), and as a key component of the response inhibition network (Verbruggen and Logan, 2008; Criaud and Boulinguez, 2013). This raises the intriguing possibility of a functional overlap between resolution of response conflict and the formation of confidence.

These roles could be unified by considering rIFG as the region in which the posterior is computed, because the posterior belief on sensory causes affords a hypothesis space for adaptive, plausible actions (Mansouri et al., 2009). Such a view is consistent with evidence for rIFG in appropriately acting on perceptual choices (Suzuki and Gottlieb, 2013), computing behavioural significance (Sakagami and Pan, 2007), computing action-outcome likelihoods that modulate motor cortex (Morris et al., 2014), and representing the posterior (d’Acremont et al., 2013). It has even been shown that the rIFG BOLD response to decision errors is associated with both the valence of the decision outcome, and the optimism of the participant (‘self-belief’; Sharot et al. 2011), consistent with a view of rIFG in which high-level, abstracted posteriors are computed from beliefs and errors. Anatomical considerations support such a view, since the rIFG is directly connected with regions relevant for both cognitive and motor control (Petrides and Pandya, 2002). We leave open for future research the question of whether and how rIFG relates perceptual confidence to action outcomes.

## Summary

In summary, we have shown that top-down expectations are integrated into decision confidence, and have shown that this occurs in a functional network consisting of rIFG, bilateral OFC and intracalcarine sulcus. Here, top-down perceptual expectations and bottom-up sensory inputs are integrated into a subjective sense of perceptual confidence. Together, our data reveal a crucial role of top-down influences in the mechanism by which perceptual decisions become available for conscious report.

## ABBREVIATED TITLE

Predictions shape confidence in rIFG

## References

Aitchison L, Bang D, Bahrami B, Latham PE (2015) Doubly Bayesian Analysis of Confidence in Perceptual Decision-Making. PLOS Comput Biol 11:e1004519 Available at: http://dx.plos.org/10.1371/journal.pcbi.1004519.

Ashburner J, Friston KJ (2000) Voxel-based morphometry–the methods. Neuroimage 11:805–821 Available at: http://www.ncbi.nlm.nih.gov/pubmed/10860804 [Accessed February 28, 2013].

Bahrami B, Olsen K, Latham PE, Roepstorff A, Rees G, Frith CD (2010) Optimally interacting minds. Science 329:1081–1085 Available at: http://www.pubmedcentral.nih.gov/articlerender.fcgi?artid=3371582&tool=pmcentrez&rendertype=abstract.

Bar M (2007) The proactive brain: using analogies and associations to generate predictions. Trends Cogn Sci 11:280–289.

Bauer M, Stenner M-P, Friston KJ, Dolan RJ (2014) Attentional Modulation of Alpha/Beta and Gamma Oscillations Reflect Functionally Distinct Processes. J Neurosci 34:16117–16125 Available at: http://www.jneurosci.org/cgi/doi/10.1523/JNEUROSCI.3474-13.2014.

Bayer HM, Glimcher PW (2005) Midbrain dopamine neurons encode a quantitative reward prediction error signal. Neuron 47:129–141.

Beck DM, Kastner S (2009) Top-down and bottom-up mechanisms in biasing competition in the human brain. Vision Res 49:1154–1165 Available at: http://www.pubmedcentral.nih.gov/articlerender.fcgi?artid=2740806&tool=pmcentrez&rendertype=abstract [Accessed August 14, 2013].

Beck DM, Rees G, Frith CD, Lavie N (2001) Neural correlates of change.:645–650.

Bitzer S, Park H, Blankenburg F, Kiebel SJ (2014) Perceptual decision making: drift-diffusion model is equivalent to a Bayesian model. Front Hum Neurosci 8:102 Available at: http://www.pubmedcentral.nih.gov/articlerender.fcgi?artid=3935359&tool=pmcentrez&rendertype=abstract [Accessed September 25, 2014].

Bubic A, von Cramon DY, Jacobsen T, Schröger E, Schubotz RI (2009) Violation of expectation: neural correlates reflect bases of prediction. J Cogn Neurosci 21:155–168 Available at: http://www.ncbi.nlm.nih.gov/pubmed/18476761.

Bülthoff I, Bülthoff H, Sinha P (1998) Top-down influences on stereoscopic depth-perception. Nat Neurosci 1:254–257.

Casey BJ, Thomas KM, Welsh TF, Badgaiyan RD, Eccard CH, Jennings JR, Crone EA (2000) Dissociation of response conflict, attentional selection, and expectancy with functional magnetic resonance imaging. Proc Natl Acad Sci 97:8728–8733.

Clark A (2013) The many faces of precision (Replies to commentaries on “Whatever next? Neural prediction, situated agents, and the future of cognitive science”). Front Psychol 4:270 Available at: http://www.pubmedcentral.nih.gov/articlerender.fcgi?artid=3659294&tool=pmcentrez&rendertype=abstract [Accessed September 16, 2013].

Clos M, Langner R, Meyer M, Oechslin MS, Zilles K, Eickhoff SB (2014) Effects of prior information on decoding degraded speech: An fMRI study. Hum Brain Mapp 35:61–74.

Criaud M, Boulinguez P (2013) Have we been asking the right questions when assessing response inhibition in go/no-go tasks with fMRI? A meta-analysis and critical review. Neurosci Biobehav Rev 37:11–23 Available at: http://www.sciencedirect.com/science/article/pii/S0149763412001935 [Accessed November 3, 2015].

d’Acremont M, Schultz W, Bossaerts P (2013) The Human Brain Encodes Event Frequencies While Forming Subjective Beliefs. J Neurosci 33:10887–10897 Available at: http://www.jneurosci.org/cgi/doi/10.1523/JNEUROSCI.5829-12.2013.

de Lange FP, Rahnev D a, Donner TH, Lau H, Lange FP De, Rahnev D a, Donner TH, Lau H (2013) Prestimulus oscillatory activity over motor cortex reflects perceptual expectations. J Neurosci 33:1400–1410 Available at: http://www.ncbi.nlm.nih.gov/pubmed/23345216 [Accessed May 29, 2014].

De Martino B, Fleming SM, Garrett N, Dolan RJ (2013) Confidence in value-based choice. Nat Neurosci 16:105–110 Available at: http://www.pubmedcentral.nih.gov/articlerender.fcgi?artid=3786394&tool=pmcentrez&rendertype=abstract [Accessed July 10, 2014].

Dienes Z (2008) Subjective measures of unconscious knowledge. Prog Brain Res 168:49–64 Available at: http://www.ncbi.nlm.nih.gov/pubmed/18166385 [Accessed November 11, 2013].

Downar J, Crawley a P, Mikulis DJ, Davis KD (2000) A multimodal cortical network for the detection of changes in the sensory environment. Nat Neurosci 3:277–283.

Downar J, Crawley AP, Mikulis DJ, Davis KD (2002) A Cortical Network Sensitive to Stimulus Salience in a Neutral Behavioral Context Across Multiple Sensory Modalities. J Neurophysiol 87:615–620 Available at: http://jn.physiology.org/content/87/1/615.short [Accessed November 20, 2015].

Egner T, Monti JM, Summerfield C (2010) Expectation and surprise determine neural population responses in the ventral visual stream. J Neurosci 30:16601–16608 Available at: http://www.ncbi.nlm.nih.gov/pubmed/21147999 [Accessed March 11, 2013].

Eickhoff SB, Stephan KE, Mohlberg H, Grefkes C, Fink GR, Amunts K, Zilles K (2005) A new SPM toolbox for combining probabilistic cytoarchitectonic maps and functional imaging data. Neuroimage 25:1325–1335.

Engel AK, Fries P, Singer W (2001) Dynamic predictions: oscillations and synchrony in top-down processing. Nat Rev Neurosci 2:704–716.

Evans S, Azzopardi P (2007) Evaluation of a “bias-free” measure of awareness. Spat Vis 20:61–77 Available at: http://www.ncbi.nlm.nih.gov/pubmed/17357716.

Fetsch C, Kiani R, Newsome W, Shadlen M (2014) Effects of Cortical Microstimulation on Confidence in a Perceptual Decision. Neuron.

Fetsch CR, Kiani R, Shadlen MN (2015) Predicting the Accuracy of a Decision: A Neural Mechanism of Confidence. Cold Spring Harb Symp Quant Biol 79:185–197 Available at: http://symposium.cshlp.org/lookup/doi/10.1101/sqb.2014.79.024893.

Fiser J, Berkes P, Orbán G, Lengyel M (2010) Statistically optimal perception and learning: from behavior to neural representations. Trends Cogn Sci 14:119–130 Available at: http://linkinghub.elsevier.com/retrieve/pii/S1364661310000045.

Fleck MS, Daselaar SM, Dobbins IG, Cabeza R (2006) Role of prefrontal and anterior cingulate regions in decision-making processes shared by memory and nonmemory tasks. Cereb Cortex 16:1623–1630 Available at: http://www.ncbi.nlm.nih.gov/pubmed/16400154 [Accessed May 22, 2015].

Fleming SM, Dolan RJ (2012) Review. Neural basis of metacognition. Philos Trans R Soc B Biol Sci 367:1338–1349.

Fleming SM, Huijgen J, Dolan RJ (2012) Prefrontal contributions to metacognition in perceptual decision making. J Neurosci 32:6117–6125 Available at: http://www.pubmedcentral.nih.gov/articlerender.fcgi?artid=3359781&tool=pmcentrez&rendertype=abstract [Accessed February 27, 2013].

Fleming SM, Lau HC (2014) How to measure metacognition. Front Hum Neurosci 8:1–9 Available at: http://www.frontiersin.org/Human_Neuroscience/10.3389/fnhum.2014.00443/abstract [Accessed July 16, 2014].

Friston K (2009) The free-energy principle: a rough guide to the brain? Trends Cogn Sci 13:293–301 Available at: http://www.ncbi.nlm.nih.gov/pubmed/19559644 [Accessed July 11, 2014].

Garrido MI, Kilner JM, Kiebel SJ, Friston KJ (2009) Dynamic causal modeling of the response to frequency deviants. J Neurophysiol 101:2620–2631.

Gherman S, Philiastides MG (2015) Neural representations of confidence emerge from the process of decision formation during perceptual choices. Neuroimage 106:134–143 Available at: http://linkinghub.elsevier.com/retrieve/pii/S1053811914009537.

Gilbert CD, Li W (2013) Top-down influences on visual processing. Nat Rev Neurosci 14:350–363 Available at: http://www.nature.com/doifinder/10.1038/nrn3476 [Accessed April 18, 2013].

Grinband J, Hirsch J, Ferrera VP (2006) A neural representation of categorization uncertainty in the human brain. Neuron 49:757–763.

Hampshire A, Chamberlain SR, Monti MM, Duncan J, Owen AM (2010) The role of the right inferior frontal gyrus: inhibition and attentional control. Neuroimage 50:1313–1319 Available at: http://www.sciencedirect.com/science/article/pii/S1053811909013986 [Accessed November 10, 2014].

Hebart MN, Schriever Y, Donner TH, Haynes J-DJ-D (2014) The Relationship between Perceptual Decision Variables and Confidence in the Human Brain. Cereb Cortex:bhu181 – Available at: http://www.ncbi.nlm.nih.gov/pubmed/25112281 [Accessed October 29, 2014].

Hilgenstock R, Weiss T, Witte OW (2014) You’d Better Think Twice: Post-Decision Perceptual Confidence. Neuroimage 99:323–331 Available at: http://dx.doi.org/10.1016/j.neuroimage.2014.05.049.

Jiang J, Summerfield C, Egner T (2013) Attention sharpens the distinction between expected and unexpected percepts in the visual brain. J Neurosci 33:18438–18447 Available at: http://www.pubmedcentral.nih.gov/articlerender.fcgi?artid=3834051&tool=pmcentrez&rendertype=abstract [Accessed May 29, 2014].

John-saaltink ES, Utzerath C, Kok P, Lau HC (2015) Expectation Suppression in Early Visual Cortex Depends on Task Set.:1–14.

Kass RE, Raftery AE (1995) Bayes Factors. J Am Stat Assoc 90:773–795.

Kepecs A, Mainen ZF (2012) A computational framework for the study of confidence in humans and animals. Philos Trans R Soc Lond B Biol Sci 367:1322–1337 Available at: http://www.ncbi.nlm.nih.gov/pubmed/22492750 [Accessed February 27, 2013].

Kepecs A, Uchida N, Zariwala HA, Mainen ZF (2008) Neural correlates, computation and behavioural impact of decision confidence. Nature 455:227–231 Available at: http://www.ncbi.nlm.nih.gov/pubmed/18690210 [Accessed February 27, 2013].

Kiani R, Corthell L, Shadlen MN (2014) Choice Certainty Is Informed by Both Evidence and Decision Time. Neuron 84:1329–1342 Available at: http://dx.doi.org/10.1016/j.neuron.2014.12.015.

Kiani R, Shadlen MN (2009) Representation of confidence associated with a decision by neurons in the parietal cortex. Science (80-) 324:759–764.

Kim H, Shimojo S, O’Doherty JP (2011) Overlapping Responses for the Expectation of Juice and Money Rewards in Human Ventromedial Prefrontal Cortex. Cereb Cortex 21:769–776 Available at: http://www.cercor.oxfordjournals.org/cgi/doi/10.1093/cercor/bhq145.

Kok P, Jehee JF, de Lange FP (2012) Less Is More: Expectation Sharpens Representations in the Primary Visual Cortex. Neuron 75:265–270.

Kok P, Rahnev D, Jehee JFM, Lau HC, de Lange FP (2011) Attention reverses the effect of prediction in silencing sensory signals. Cereb Cortex 22:2197–2206 Available at: http://www.ncbi.nlm.nih.gov/pubmed/22047964 [Accessed March 5, 2013].

Kouider S, Long B, Le Stanc L, Charron S, Fievet A-C, Barbosa LS, Gelskov S V (2015) Neural dynamics of prediction and surprise in infants. Nat Commun 6 Available at: http://dx.doi.org/10.1038/ncomms9537.

Larsson J, Smith AT (2012) fMRI repetition suppression: neuronal adaptation or stimulus expectation? Cereb Cortex 22:567–576 Available at: http://www.pubmedcentral.nih.gov/articlerender.fcgi?artid=3278317&tool=pmcentrez&rendertype=abstract [Accessed May 29, 2014].

Law JR, Flanery M a, Wirth S, Yanike M, Smith AC, Frank LM, Suzuki W a, Brown EN, Stark CEL (2005) Functional magnetic resonance imaging activity during the gradual acquisition and expression of paired-associate memory. J Neurosci 25:5720–5729.

Lebreton M, Abitbol R, Daunizeau J, Pessiglione M (2015) Automatic integration of confidence in the brain valuation signal. Nat Neurosci 18 Available at: http://www.ncbi.nlm.nih.gov/pubmed/26192748 [Accessed July 24, 2015].

Lee TS (2002) Top-down influence in early visual processing: a Bayesian perspective. Physiol Behav 77:645–650.

Love, J., Selker, R., Marsman, M., Jamil, T., Dropmann, D., Verhagen, A. J., Ly, A., Gronau, Q. F., Smira, M., Epskamp, S., Matzke, D., Wild, A., Rouder, J. N., Morey, R. D. & Wagenmakers E-J (2015) JASP (Version 0.7).

Maloney LT, Dal Martello MF, Sahm C, Spillmann L (2005) Past trials influence perception of ambiguous motion quartets through pattern completion. Proc Natl Acad Sci U S A 102:3164–3169.

Mansouri F a, Tanaka K, Buckley MJ (2009) Conflict-induced behavioural adjustment: a clue to the executive functions of the prefrontal cortex. Nat Rev Neurosci 10:141–152 Available at: http://www.ncbi.nlm.nih.gov/pubmed/19153577 [Accessed May 20, 2015].

Meyniel F, Schlunegger D, Dehaene S (2015a) The Sense of Confidence during Probabilistic Learning: A Normative Account. PLoS Comput Biol 11:e1004305 Available at: http://www.ncbi.nlm.nih.gov/pubmed/26076466.

Meyniel F, Sigman M, Mainen ZF (2015b) Confidence as Bayesian Probability: From Neural Origins to Behavior. Neuron 88:78–92.

Morales J, Solovey G, Maniscalco B, Rahnev D, de Lange FP, Lau H (2015) Low attention impairs optimal incorporation of prior knowledge in perceptual decisions. Atten Percept Psychophys 77:2021–2036 Available at: http://www.ncbi.nlm.nih.gov/pubmed/25836765 [Accessed August 28, 2015].

Morris RW, Dezfouli A, Griffiths KR, Balleine BW (2014) Action-value comparisons in the dorsolateral prefrontal cortex control choice between goal-directed actions. Nat Commun 5:4390 Available at: http://www.pubmedcentral.nih.gov/articlerender.fcgi?artid=4124863&tool=pmcentrez&rendertype=abstract [Accessed May 22, 2015].

Nakahara H, Itoh H, Kawagoe R, Takikawa Y, Hikosaka O (2004) Dopamine Neurons Can Represent Context-Dependent Prediction Error. Neuron 41:269–280.

Nassar MR, Wilson RC, Heasly B, Gold JI (2010) An Approximately Bayesian Delta-Rule Model Explains the Dynamics of Belief Updating in a Changing Environment. J Neurosci 30:12366–12378 Available at: http://www.jneurosci.org/cgi/doi/10.1523/JNEUROSCI.0822-10.2010.

Overgaard M, Sandberg K (2012) Kinds of access: different methods for report reveal different kinds of metacognitive access. Philos Trans R Soc Lond B Biol Sci 367:1287–1296 Available at: http://www.ncbi.nlm.nih.gov/pubmed/22492747 [Accessed March 2, 2013].

Pajani a., Kok P, Kouider S, de Lange FP (2015) Spontaneous Activity Patterns in Primary Visual Cortex Predispose to Visual Hallucinations. J Neurosci 35:12947–12953 Available at: http://www.jneurosci.org/cgi/doi/10.1523/JNEUROSCI.1520-15.2015.

Petrides M, Pandya DN (2002) Comparative cytoarchitectonic analysis of the human and the macaque ventrolateral prefrontal cortex and corticocortical connection patterns in the monkey. Eur J Neurosci 16:291–310 Available at: http://doi.wiley.com/10.1046/j.1460-9568.2001.02090.x [Accessed November 3, 2015].

Petrusic WM, Baranski J V (2003) Judging confidence influences decision processing in comparative judgments. Psychon Bull Rev 10:177–183.

Pleskac TJ, Busemeyer JR (2011) Two-stage dynamic signal detection: A theory of choice, decision time, and confidence. Psychol Rev 118:56.

Rahnev D, Maniscalco B, Graves T, Huang E, de Lange FP, Lau H (2011) Attention induces conservative subjective biases in visual perception. Nat Neurosci 14:1513–1515 Available at: http://www.ncbi.nlm.nih.gov/pubmed/22019729 [Accessed September 20, 2013].

Ratcliff R, Starns JJ (2009) Modelling confidence and response time in recognition memory. Psychol Rev 116:59–83.

Rorden C, Brett M (2000) Stereotaxic display of brain lesions. Behav Neurol 12:191–200.

Sakagami M, Pan X (2007) Functional role of the ventrolateral prefrontal cortex in decision making. Curr Opin Neurobiol 17:228–233.

Sandberg K, Timmermans B, Overgaard M, Cleeremans A (2010) Measuring consciousness: is one measure better than the other? Conscious Cogn 19:1069–1078 Available at: http://dx.doi.org/10.1016/j.concog.2009.12.013 [Accessed November 11, 2013].

Schoenbaum G, Roesch M (2005) Orbitofrontal cortex, associative learning, and expectancies. Neuron 47:633–636 Available at: http://www.ncbi.nlm.nih.gov/pubmed/16129393.

Seriès P, Seitz AR (2013) Learning what to expect (in visual perception). Front Hum Neurosci 7:1–14 Available at: http://journal.frontiersin.org/article/10.3389/fnhum.2013.00668/abstract.

Seth AK, Dienes Z, Cleeremans A, Overgaard M, Pessoa L (2008) Measuring consciousness: relating behavioural and neurophysiological approaches. Trends Cogn Sci 12:314–321 Available at: http://www.pubmedcentral.nih.gov/articlerender.fcgi?artid=2767381&tool=pmcentrez&rendertype=abstract [Accessed February 28, 2013].

Sharot T, Korn CW, Dolan RJ (2011) How unrealistic optimism is maintained in the face of reality. Nat Neurosci 14:1475–1479 Available at: http://dx.doi.org/10.1038/nn.2949.

Sherman MT, Seth AK, Barrett AB, Kanai R (2015) Prior expectations facilitate metacognition for perceptual decision. Conscious Cogn 35:53–65 Available at: http://www.sciencedirect.com/science/article/pii/S1053810015000926.

Smith FW, Muckli L (2010) Nonstimulated early visual areas carry information about surrounding context. Proc Natl Acad Sci U S A 107:20099–20103.

Stalnaker T, Cooch NK, Schoenbaum G (2015) What the orbitofrontal cortex does not do. Nat Neurosci 18:620–627.

Suzuki M, Gottlieb J (2013) Distinct neural mechanisms of distractor suppression in the frontal and parietal lobe. Nat Neurosci 16:98–104 Available at: http://www.pubmedcentral.nih.gov/articlerender.fcgi?artid=4207121&tool=pmcentrez&rendertype=abstract [Accessed April 23, 2015].

Trapp S, Bar M (2015) Prediction, context, and competition in visual recognition. Ann N Y Acad Sci 1339:190–198 Available at: http://doi.wiley.com/10.1111/nyas.12680.

Verbruggen F, Logan GD (2008) Response inhibition in the stop-signal paradigm. Trends Cogn Sci 12:418–424 Available at: http://www.pubmedcentral.nih.gov/articlerender.fcgi?artid=2709177&tool=pmcentrez&rendertype=abstract [Accessed June 19, 2015].

Wacongne C, Labyt E, Van Wassenhove V, Bekinschtein T, Naccache L, Dehaene S (2011) Evidence for a hierarchy of predictions and prediction errors in human cortex. Proc Natl Acad Sci 108:1–6 Available at: http://www.pnas.org/cgi/doi/10.1073/pnas.1117807108 [Accessed March 7, 2013].

Wallis JD (2007) Orbitofrontal cortex and its contribution to decision-making. Annu Rev Neurosci 30:31–56 Available at: http://www.ncbi.nlm.nih.gov/pubmed/17417936 [Accessed February 3, 2015].

Wierzchoń M, Paulewicz B, Asanowicz D, Timmermans B, Cleeremans A (2014) Different subjective awareness measures demonstrate the influence of visual identification on perceptual awareness ratings. Conscious Cogn 27C:109–120 Available at: http://www.ncbi.nlm.nih.gov/pubmed/24842312 [Accessed July 14, 2014].

Yeung N, Summerfield C (2012) Metacognition in human decision-making: confidence and error monitoring. Philos Trans R Soc Lond B Biol Sci 367:1310–1321 Available at: http://www.pubmedcentral.nih.gov/articlerender.fcgi?artid=3318764&tool=pmcentrez&rendertype=abstract [Accessed March 2, 2013].

Yuille A, Kersten D (2006) Vision as Bayesian inference: analysis by synthesis? Trends Cogn Sci 10:301–308.

